# Genetics-driven Risk Predictions with Differentiable Mendelian Randomization

**DOI:** 10.1101/2024.03.06.583727

**Authors:** Daniel Sens, Ludwig Gräf, Liubov Shilova, Francesco Paolo Casale

## Abstract

Accurate predictive models of future disease onset are crucial for effective preventive healthcare, yet longitudinal datasets linking early risk factors to subsequent health outcomes are scarce. To address this challenge, we introduce Differentiable Mendelian Randomization (DMR), an extension of the classical Mendelian Randomization framework to learn risk predictors without longitudinal data. To do so, DMR leverages risk factors and genetic data from a healthy cohort, along with results from genome-wide association studies (GWAS) of diseases of interest. After training, the learned predictor can be used to assess risk for new patients solely based on risk factors. We validated DMR through comprehensive simulations and in future type 2 diabetes predictions in UK Biobank participants without diabetes, using follow-up onset labels for validation. Finally, we apply DMR to predict future Alzheimer’s onset from brain imaging biomarkers. Overall, with DMR we offer a new perspective in predictive modeling, showing it is possible to learn risk predictors leveraging genetics rather than longitudinal data.

## Introduction

Large biobanks such as UK Biobank (UKB) [1], the German National Cohort [2], and others [3– 6], have unlocked access to extensive health metrics and risk factors for healthy populations, enabling future disease onset predictions for prevention. Yet, the scarcity of complete follow-ups and the presence of missing data can complicate the study of disease onset [7], particularly for less prevalent diseases.

Mendelian Randomization (MR) is pivotal for identifying causal links between risk factors and health outcomes, utilizing genetic data across multiple cohorts [8,9]. For instance, MR has elucidated the causal impact of risk factors such as cholesterol levels on cardiovascular disease, invalidating the protective role of HDL cholesterol [10] and confirming the adverse effects of LDL [11]. As interest grows in using MR for preventive healthcare [12–16], we explore its potential for disease risk predictions as an alternative to longitudinal studies.

Expanding on the MR framework, we present Differentiable Mendelian Randomization (DMR), an extension for learning disease risk predictors through nonlinear functions of multiple risk factors. To achieve this, DMR utilizes risk factors and genetic data from a healthy cohort (**Figure 1a**), and external disease-specific GWAS results (**Figure 1b**). During training, DMR fine-tunes the predictor to ensure that the genetic effects on both the predictor and the disease outcome are aligned across selected genetic variants (**Figure 1c**), upholding MR’s foundational principles. Once trained, disease risk in new patients can be assessed solely using risk factors (**Figure 1d**).

**Figure 1.**
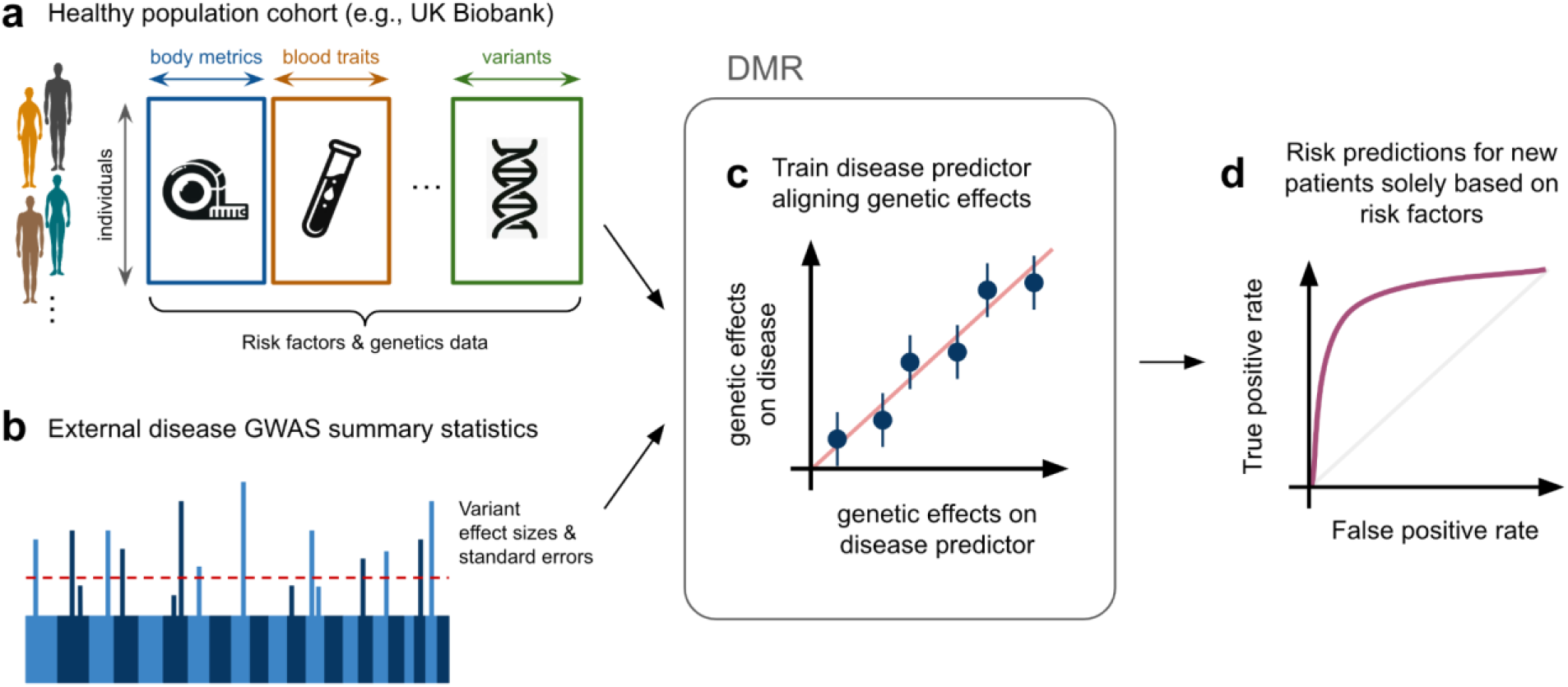
Overview of the DMR Framework for Disease Risk Prediction. (**a**) DMR utilizes health metrics and genetic data from a cohort of healthy individuals. (**b**) It integrates these with disease-specific GWAS summary statistics from an external cohort. (**c**) The framework trains risk predictors to align genetic effects with those observed in disease outcomes, maintaining adherence to classic MR principles. (**d**) Post-training, the model’s accuracy in predicting disease risk is evaluated, for example, through the receiver operating characteristic curve against actual follow-up disease onset data.

We validate DMR’s risk predictions through extensive simulations and in a type 2 diabetes (T2D) prediction task, leveraging follow-up labels for validation. Finally, we apply DMR to identify brain imaging predictors of future AD onset.

## Results

### Risk predictions with Two-sample Mendelian randomization

Two-sample Mendelian randomization (MR) utilizes GWAS summary statistics of a risk factor (exposure; e.g., LDL cholesterol) and a disease (outcome; e.g., cardiovascular disease) from different cohorts to assess the causal impact of the exposure on the outcome (**Figure 2a**). Intuitively, MR operates under the premise that, given essential assumptions, if an exposure causally influences an outcome, then the effects of exposure-associated variants on the outcome should be directly proportional to their effects on the exposure, with the slope of this proportionality quantifying the causal effect [9]. Technically, for *S* independent exposure-associated variants, the causal effect 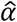 is estimated through Inverse Variance Weighting (IVW) regression [17], where the genetic effects on the outcome (*β*_*o*_ *∈R*^*S*^) are regressed on the genetic effects on the exposure (*β*_*e*_ *∈R*^*S*^), while accounting for their standard errors (*s*_*o*_ *∈R*^*S*^) (**Figure 2b**). A critical and challenging assumption of MR is the absence of horizontal pleiotropy; that is, exposure-associated genetic variants must affect the outcome solely via the exposure, without alternative pathways [18].

**Figure 2.**
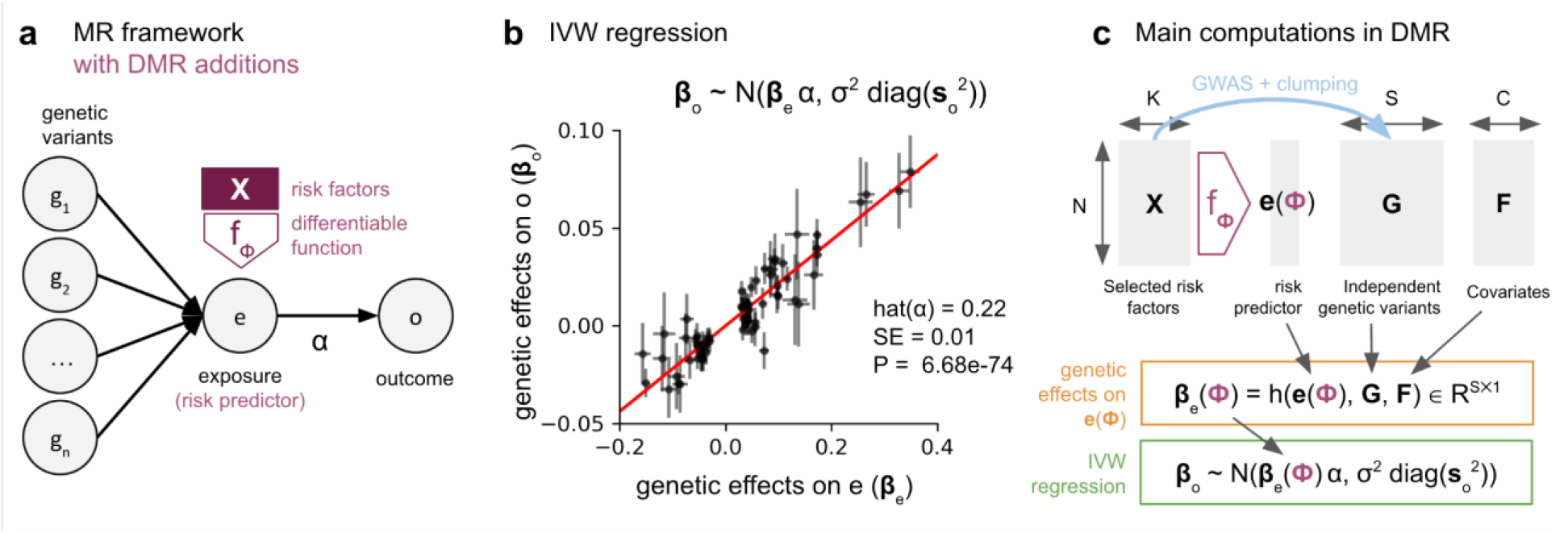
Overview of the Differentiable Mendelian Randomization framework. **(a)** Diagram illustrating MR causal assumptions, where genetic variants (*g*_*1*_, *…*, *g*_*S*_) influence exposure (e), which in turn affects outcome (o) with causal effect *α*. Additions unique to DMR are highlighted in purple: the exposure is a risk predictor, which is computed as a differentiable function *f*_*ϕ*_ (parametrized by *ϕ*) of risk factors *X*. (**b**) Illustration of IVW regression, where genetic variant effects on outcome (*β*_*o*_) are regressed against their impact on exposure (*β*_*e*_), accounting for their standard errors (*s*_*o*_). (**c**) Main computations in DMR, including computation of the risk predictor *e(ϕ)*, the estimation of genetic effects on the risk predictor *β*_*e*_*(ϕ)*, and the computation of the IVW regression loss. The function *h(e(ϕ), G, F)* returns marginal regression weights of each variant *G*_*:s*_ on *e(ϕ)* accounting for covariates *F*. As all these steps are differentiable, *f*_*ϕ*_ can be learned through gradient-based optimization of the IVW regression loss.

When both early risk factors and follow-up disease labels are available, conventional supervised learning techniques can model future disease onset *y* as a function *f(x)* of potential risk factors **x** *∈ℝ*^***K***^ through classical supervised learning. Yet, when follow-up data is scarce, Mendelian Randomization (MR) can provide a viable alternative for constructing risk predictors *f(x)* using disease-specific GWAS statistics from an external cohort. For example, we can use MR to determine which of the *K* candidate risk factors causally affect the disease using GWAS summary statistics [16], and construct a linear risk predictor 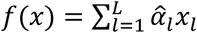 from *L* significant causal factors, where 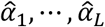 represent causal effect estimates derived by MR. Upon this basis, we extend the MR framework to support end-to-end learning of disease risk predictors.

### Differentiable Mendelian Randomization

DMR can learn risk predictors as nonlinear functions of multiple risk factors, leveraging individual-level data from a genetic cohort of healthy individuals and disease-specific GWAS summary statistics. To do so, it introduces a differentiable function *f* parametrized by *ϕ* within the MR framework to aggregate multiple risk factors into a single risk predictor (**Figure 2a**). The predictor is then fine tuned to optimize the IVW regression. Briefly, for *N* individuals, *K* risk factors *X ∈R*^*N*×*K*^, *C* covariates *F ∈R*^*N*×*C*^ and *S* independent genetic variants *G ∈R*^*N*×*S*^ associated with at least one of the *K* risk factor, the IVW regression loss can be computed as follows (**Figure 2c**):

1. Compute aggregate risk predictor *e(ϕ) ∈R*^*N*×*1*^ from *X* using *f*_*ϕ*_;
2. Compute genetic effects *β*_*e*_*(ϕ) ∈R*^*S*×*1*^ on the aggregate risk predictor as the marginal regression weights of each variant *G*_*:s*_ on *e(ϕ)* accounting for covariates *F*. This step mirrors the exposure GWAS step in standard MR;
3. Compute IVW regression loss based on risk predictor genetic effects *β*_*e*_*(ϕ)*, and outcome statistics *β*_*o*_ and *s*_*o*_, i.e. *L*_*IVW*_*(ϕ, α, σ*^*2*^*) = −log𝒩( β*_*o*_ ∣ *β*_*e*_*(ϕ) α, σ*^*2*^ *diag(s*_*o*_)).

As all these steps are differentiable, *ϕ* can be learned through gradient-based optimization of the IVW loss (**Methods**). To select independent risk factor-associated variants for our analyses, we performed univariate GWAS analyses for each risk factor followed by a multivariate clumping procedure (**Methods**). We considered the following nonlinear function of risk factors:

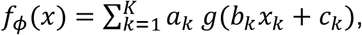

with parameters *ϕ = {a*_*1*_, *…*, *a*_*K*_, *b*_*1*_, *…*, *b*_*K*_, *c*_*1*_, *…*, *c*_*K*_*}*, where *g* is a J-shaped warping function, a choice that enables modeling potential nonlinearities while being simple and clinically plausible. Such a shape function is commonly used in risk prediction as it captures the scenario where contributions from single risk factors remain minimal until a critical threshold and then escalate [19–21].

### Validation of DMR using simulated data

We evaluated the proposed DMR framework through a series of simulations derived from the UKB, encompassing 309,846 unrelated European individuals. We focused on 26 blood traits as potential risk factors, simulating scenarios where subsets of these traits have a causal influence on the simulated health outcomes. Importantly, the health outcome was modeled as a linear combination of these traits, each transformed by a J-shaped function to represent contributions that activate beyond specific thresholds (**Methods**). Our simulation framework allowed us to examine the efficacy of DMR under various conditions, including the presence of horizontal pleiotropy and varying degrees of risk factor influence on outcome variance.

We compared DMR with its linear variant (DMR-LIN) and linear risk predictors informed by two-sample univariate and multivariable Mendelian Randomization (UVMR and MVMR, respectively; **Methods**). Beyond MR-derived models, we included a longitudinal reference model (LRM) trained directly on individual-level follow-up labels as a performance benchmark (**Methods**). Our evaluation maintained a strict two-sample framework, preventing any overlap between the cohorts used for determining genetic effects on risk factors and outcomes. We measured the accuracy of risk predictions for all methods using the Spearman correlation, comparing estimated risk scores versus simulated ones in a held-out validation set. To ensure calibration of our evaluation procedure, we verified models’ performance was equivalent to random chance in control simulations without causal links (**Supplementary Figure A1**).

The findings from our simulations underscore the robustness and versatility of DMR across a wide range of scenarios. Specifically, DMR’s performance in estimating risk remained stable when increasing the number of causal risk factors (**Figure 3a**). This superior performance persisted across different values of the variance explained by the risk factors (**Figure 3b**) and when simulating horizontal pleiotropy (**Figure 3c**; **Methods**). Finally, we note that although the longitudinal reference model offers the highest predictive accuracy, DMR’s performance can be competitive if follow-up data is sparse (**Supplementary Figure A2**). Collectively, these results highlight DMR’s potential as a powerful tool for predictive modeling when follow-up labels are scarce.

**Figure 3.**
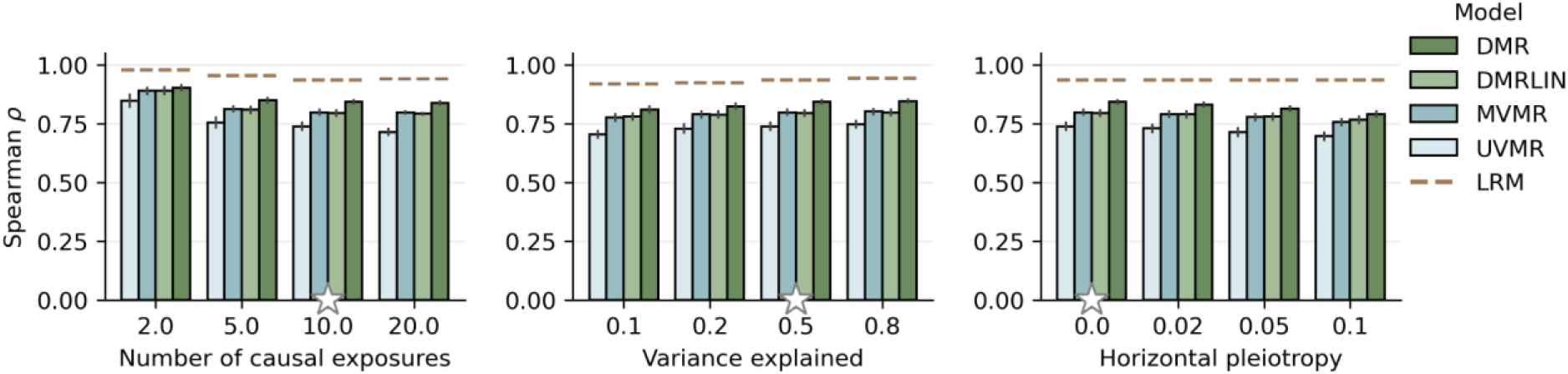
Assessment of disease risk prediction accuracy using simulated data. Comparison of model accuracy in recovering the simulated exposure measured by Spearman correlation. Compared are DMR, its linear variant (DMR-LIN), multivariable MR (MVMR), univariate MR (UVMR), and the supervised model accessing individual-level follow-up labels (LRM; **Methods**), varying the number of simulated causal exposures (**a**), the fraction of outcome variance explained by the risk factors (**b**), and the fraction of outcome variance explained by horizontal pleiotropy (**c**). Stars denote standard values held constant while other parameters were varied. Error bars indicate standard errors across 10 replicate experiments.

### Validation of DMR in Predicting 5-Year Type 2 Diabetes Risk

Next, we evaluated the prediction accuracy of DMR in a real-world setting, considering a type 2 diabetes (T2D) dataset derived from the UKB dataset. Specifically, we aimed to predict 5-year T2D risk by leveraging risk factors and genetic data from 218,665 UKB individuals with no reported T2D at the initial assessment (**Methods**), and external GWAS summary statistics for T2D [22]. As input risk factors, we used 37 traits previously linked to diabetes risk [23], including metabolic, anthropometric, and cardiovascular metrics (**Figure 4a**). As genetic instruments, we used 6,077 independent genetic variants associated with at least one of the risk traits at the genome-wide significance level (*P* < 5 ⋅ *10*^*−8*^; **Methods**).

**Figure 4.**
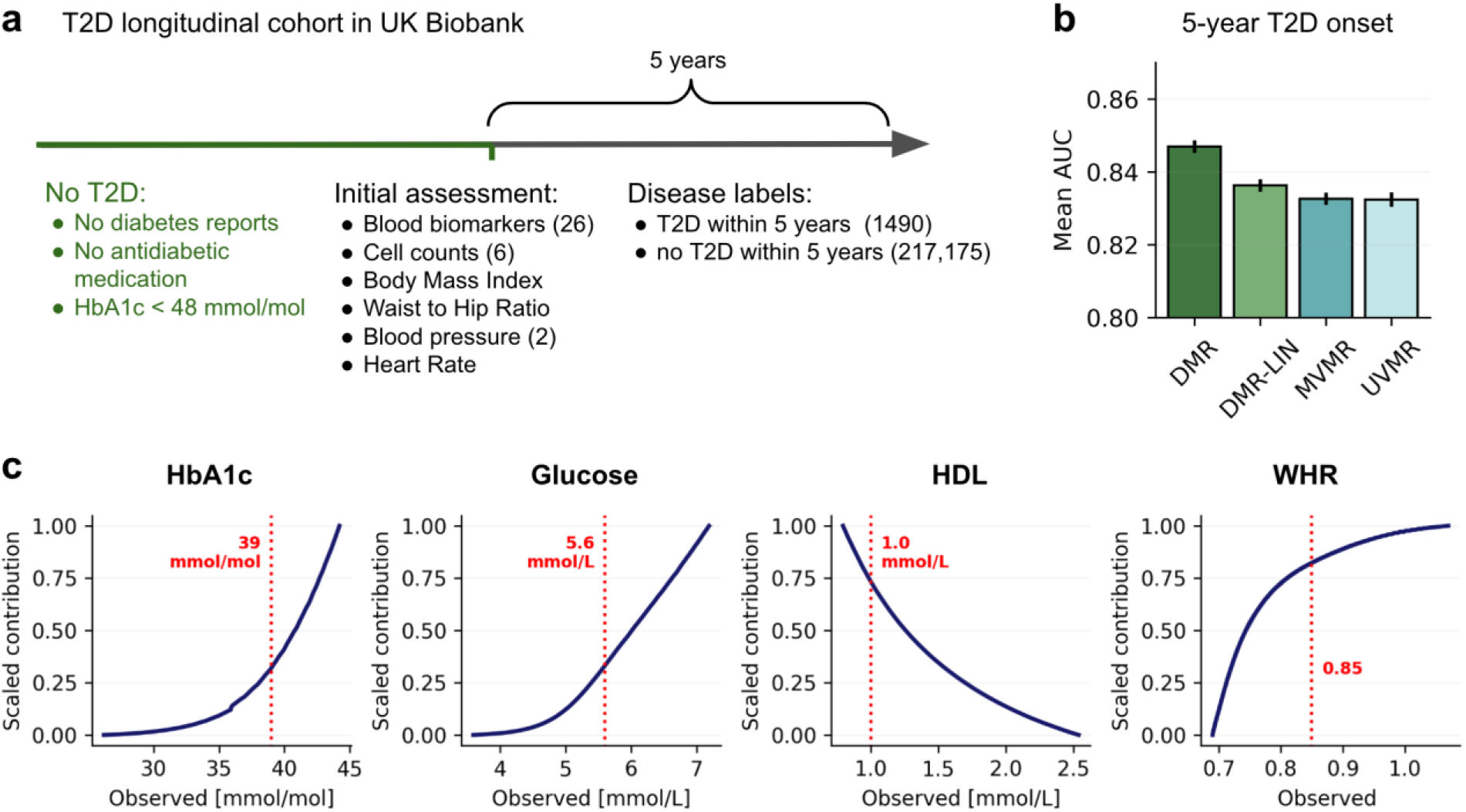
Validation of DMR in Predicting 5-Year T2D Risk. (**a**) Schematic representation of UKB T2D cohort, showing the inclusion criteria, the 37 risk factors included in our analysis, and the definition of the 5-year T2D onset labels. (**b**) Comparison of the mean Area Under the receiver operating characteristic Curve (AUC) scores for 5-year T2D onset labels obtained using DMR, its linear variant (DMR-LIN), multivariable MR (MVMR) and univariate MR (UVMR). Error bars denote standard errors across 50 random train/test splits (**Methods**). (**c**) Scaled contributions to risk learned by DMR as function of observed values for glycated hemoglobin (HbA1c), glucose, high-density lipoprotein (HDL) and waist-to-hip ratio (WHR). Risk reference thresholds are annotated in red.

DMR outperformed baseline MR methods, achieving an average Area Under the receiver operating characteristic Curve (AUC) of 0.847 (± 0.002) against 0.836 (± 0.002) obtained using MVMR (P < 10^−4^; **Figure 4b, Supplementary Figure A3**). Additionally, we evaluated MR-based model predictions against a polygenic risk score (PRS) model that relies exclusively on genetic data for predictions [24] (**Methods**), unlike MR-based models which use risk factors for prediction. Notably, the PRS model markedly underperformed compared to MR-based models, recording an AUC of 0.647 (±0.002) (**Supplementary Figure A3**). Although a supervised reference model expectedly yielded the best performance when trained on full individual-level follow-up data, DMR demonstrated competitiveness in scenarios with low numbers of follow-up labels (**Supplementary Figure A2**).

The risk predictor derived from DMR robustly aligns with established clinical knowledge, underscored by its correlation with individual factors (**Supplementary Figure A4**) [23]. Notably, the nonlinear relationships identified by DMR align with clinical expectations; for example, the risk contributions from glycated hemoglobin and glucose show significant increases nearing clinical risk thresholds (**Figure 4c**). Overall, these results showcase the accuracy and clinical plausibility of risk predictors learned through DMR in a real data setting.

### Application of DMR to Predict 5-Year Alzheimer’s Disease Risk from Brain Imaging Biomarkers

We applied DMR to identify imaging biomarkers predictive of 5-year AD risk focusing on 31,552 unrelated European individuals in the UKB with brain imaging data. As imaging risk factors, we considered 70 subcortical and gray matter volumes traits from T1 MRI having at least five independent genome-wide significant signals, for a total of 353 independent genetic variants associated with at least one of these traits. As external AD GWAS results, we used AD GWAS summary statistics from [25].

Remarkably, all multivariable MR models exceeded the performance expected by chance (**Figure 5a**), with DMR achieving markedly higher accuracy compared to linear counterparts (DMR AUC at 0.741 ± 0.003 vs. DMR-LIN at 0.690 ± 0.003 vs. MVMR at 0.629 ± 0.004; **Figure 5a**). A thorough analysis of the key imaging features pinpointed by DMR for AD predictions underscored their correlation with reductions in gray matter and subcortical volume across various regions (**Supplementary Figure 5**), particularly in the midbrain (**Figure 5b**). The most affected regions, including the amygdala, thalamus, hippocampus, ventral striatum, frontal orbital cortex, and putamen (**Figure 5c**), are consistent with known AD pathology [26], underscoring the clinical relevance of these findings. Overall, these results underscore DMR’s effectiveness in utilizing genetic data for accurate risk prediction in the context of diseases with lower prevalence, such as AD.

**Figure 5.**
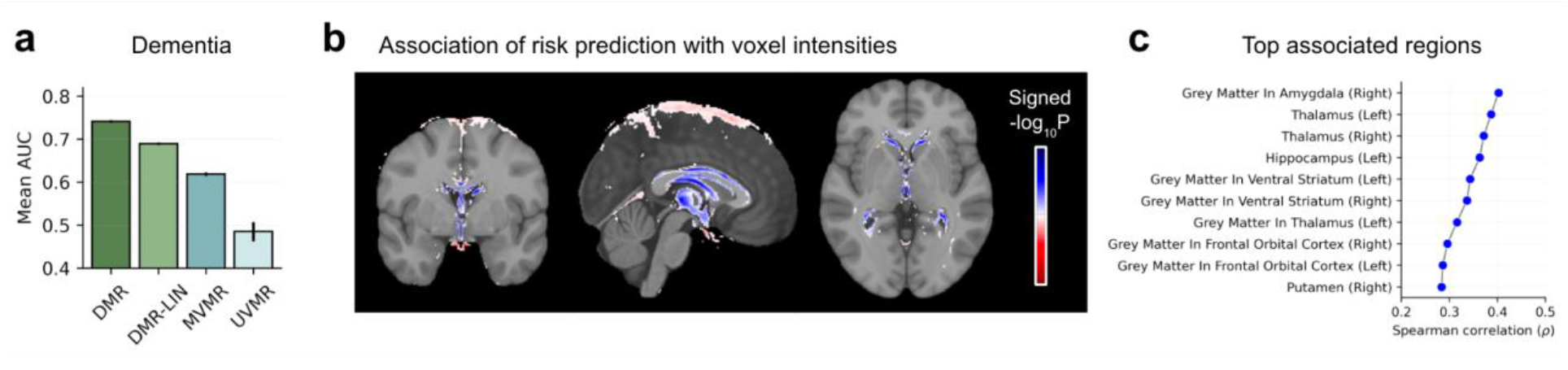
Application of DMR to Predict 5-Year AD Risk. (**a**) Comparative performance of DMR against baseline MR models using average AUC for 5-year AD predictions using follow-up labels. (**b**) Heatmap of the signed -log10 P value of association between voxel intensities and the AD risk predictor scores, overlayed on the MNI152 template [50–53]. Areas where increased risk predictor scores correlate with significant increased (decreased) voxel intensities are highlighted in red (blue) (Bonferroni-adjusted P < 0.05). (**c**) Spearman correlation coefficients between the AD risk predictor and individual MRI traits in the validation set. Results for the top 10 associated regions are displayed, with associations for all analyzed regions available in **Supplementary Figure A5**.

## Discussion

In this study, we demonstrate that the two-sample Mendelian randomization framework can enable disease risk predictions using genetic information, without relying on longitudinal data. This approach is especially valuable given that genetic biobanks boast extensive health metrics but frequently lack longitudinal disease onset data—a challenge accentuated in diseases with lower incidence rates. Specifically, we illustrate how, in such scenarios, results from large genome-wide association studies (GWAS) can serve as essential supervisory signals for training risk predictors. These signals can potentially replace or enhance longitudinal data analyses.

We extend this methodology through Differentiable MR (DMR), introducing nonlinear disease risk prediction within the MR framework. DMR’s validity is confirmed in both simulations and a longitudinal diabetes dataset, using follow-up data as benchmarks. Moreover, DMR’s application to brain imaging predictors of future AD showcases utility for less prevalent diseases. For interpretability, DMR models disease risk as a linear function of transformed risk factors, identifying risk escalations beyond specific thresholds. This feature allowed us to verify that DMR learns plausible nonlinearities for diabetes biomarkers. DMR’s flexibility, facilitated by its PyTorch implementation, promises broad applicability, from trait embeddings derived from unsupervised models [27,28] to focused biomarker discovery, such as to define new anthropometric biomarkers [29] or blood clinical scores [30].

Despite these advantages, DMR is not without limitations. The framework does not incorporate explicit mechanisms to counter weak instrument bias [31] and horizontal pleiotropy [9]— highlighting critical areas for further development. DMR requires genetic cohorts for outcome and risk factors to be sampled from the same population. Failure to meet these criteria may lead to challenges due to variations in linkage disequilibrium patterns [32] and gene-environment interactions, necessitating the integration of robust instrument selection strategies, such as variant fine-mapping [33]. Furthermore, identifying the optimal selection of genetic instruments for nonlinear MR is crucial, underscoring the need for continued research.

Although our focus has been on predicting disease risk without reliance on longitudinal data, utilizing a causal inference framework for disease prediction may provide a viable method to mitigate confounding in longitudinal datasets [7], potentially enhancing the generalization of risk predictors across different datasets. The potential of DMR to facilitate this exploration opens exciting avenues for future research, particularly as more cohorts with deeper phenotype data become available.

## Methods

### Differentiable Mendelian Randomization

#### Two-sample Mendelian Randomization and Inverse Variance Weighting

Two-sample Mendelian Randomization leverages summary statistics of GWAS of a risk factor (exposure) and a health outcome to infer the causal effect of the exposure on the outcome. Briefly, assuming *S* independent genetic variants associated with the analyzed exposure, this can be estimated through the Inverse Variance Weighting (IVW) regression [17]:

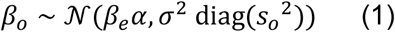

where 𝒩 denotes the multivariate normal distribution, *β*_*o*_ *∈R*^*S*^ denotes the variant effects on the outcome, *s*_*o*_ *∈R*^*S*^ their standard errors, *β*_*e*_ *∈R*^*S*^ the variant effects on the exposure, *α* the regression slope and *σ*^*2*^ the variance of the regression error. Within this framework, the causal effect of the exposure on the outcome, is the maximum likelihood estimator of the regression slope *α*, i.e., 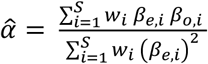 with standard error 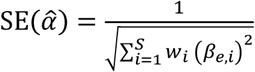, where 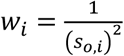. MR is an instrumental variable analysis method [34], using the genetic variants associated with the exposure as instruments. As such, it relies on key assumptions [9]: 1) The chosen instruments are robustly associated with the exposure, 2) The instruments are independent of any confounders that may influence both the exposure and the outcome, and 3) The instruments influence the outcome only through the exposure, i.e., no horizontal pleiotropy. Moreover, for the causal effect estimate to be valid, the exposure and outcome statistics need to be estimated on independent cohorts sampled from the same population [35].

#### Differentiable Mendelian Randomization

In classical two-sample MR, *β*_*e*_ is retrieved from the GWAS of a single risk factor (see Eq (1)). However, given access to individual-level data and multiple risk factors in the exposure cohort, it is feasible to define an aggregate exposure through a function of these factors and compute *β*_*e*_ by regressing genetic instruments against this aggregate exposure. Importantly, if the function *f*_*ϕ*_ parametrized by *ϕ*, is differentiable, the corresponding genetic effects on the aggregate exposure *β*_*e*_*(ϕ)* are also differentiable, enabling the learning of an aggregate risk function *f*_*ϕ*_ by directly optimizing the IVW regression loss through gradient descent. For *N* individuals, *K* risk factors *X ∈R*^*N*×*K*^, *C* covariates *F ∈R*^*N*×*C*^ and *S* independent genetic variants *G ∈R*^*N*×*S*^, each associated with at least one of the *K* risk factors, the IVW regression loss can be computed as

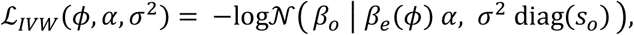

Where the genetic effects of the aggregated risk factor *β*_*e*_*(ϕ)* are computed as the marginal regression weights of each variant *G*_:,*1*_, …, *G*_:,*S*_ on *f*_*ϕ*_*(X)* accounting for covariates *F* (**Supplementary Information**). As *L*_*IVW*_ is fully differentiable in *ϕ, α, σ*^*2*^, the predictor function *f*_*ϕ*_ is end-to-end trainable. Regarding the analytical form of *f*, we opted for a linear combination of warping functions for single exposures:

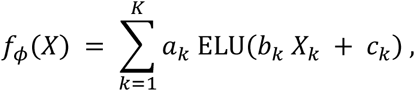

with parameters *ϕ = {a*_*1*_, *…*, *a*_*K*_, *b*_*1*_, *…*, *b*_*K*_, *c*_*1*_, *…*, *c*_*K*_*}* and where *ELU(*⋅) is the Exponential Linear Unit function [36]. This formulation assumes contributions from single risk factors remain minimal until a critical threshold is reached, after which they escalate [19–21]. Notably, by selecting a linear function for *f*_*ϕ*_, we demonstrate that DMR seamlessly reduces to multivariable Mendelian Randomization (**Supplementary Information**) [37,38], underscoring the robust foundation and adaptability of our approach. An overview of related methods to DMR is detailed in (S**upplementary Information)**

#### Bayesian model and optimization

To enhance DMR’s robustness in scenarios with a limited number of genetic instruments, we implemented a Bayesian inference approach. This involved introducing priors over the parameters *ϕ* and optimizing the log marginal likelihood of the IVW model. For parameters where analytical integration was infeasible, mean-field variational inference was utilized to derive the Evidence Lower BOund (ELBO). Optimization of the ELBO was achieved through gradient descent using the Adam optimizer, incorporating the reparameterization trick to enable backpropagation through the expectation term of the ELBO. This approach aligns with standard practices in variational inference methods that leverage gradient descent [39–41]. The learning rate for all experiments was fixed at 0.01, and we consistently applied gradient clipping with a norm bound of one while training for 1000 epochs. Risk predictions were obtained as the mean of the variational posterior of the model. Comprehensive details on our Bayesian model and variational inference procedure can be found in **Supplementary Information**. Finally, we note that prior to all experiments, risk factors were normalized using a rank-inverse Gaussian transformation, widely used phenotype transformation for GWAS analyses [42]. Our DMR framework was implemented in PyTorch [43].

#### Selection of instruments

To identify genetic variants associated with risk factors, we first conducted a univariate GWAS for each risk factor followed by a multivariate clumping procedure. GWAS analysis utilized linear regression via GCTA (fastGWA-lr functionality) [44], adjusting for sex, age, UKB array type and the top 20 genetic principal components. Adjusting for the top 20 genetic principal components is a standard practice to correct for population structure in genetic analyses of unrelated Europeans [45]. After GWAS, we applied clumping on the minimum P value statistics across all traits using PLINK [46], with parameters fixed to a P-value threshold of 5 ⋅ *10*^*−8*^, an *r*^*2*^ linkage disequilibrium cut off of *0*.*0*5, and a clumping window of 5*000* Kb, following [47]. This procedure ensured that selected variants are approximately independent and associated with at least one of the risk factors at genome-wide significant level (*P* < 5 ⋅ *10*^*−8*^).

#### Comparison models

In our study, we assess the performance of DMR in comparison to two MR-based predictive models: Univariate Mendelian Randomization (UVMR) and Multivariate Mendelian Randomization (MVMR). Both UVMR and MVMR apply a linear risk prediction function 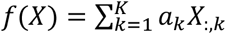, where *X* represents risk factors and *a*_*k*_ the estimated causal effects. UVMR determines *a*_*k*_ from MR-estimated causal effects 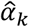 for each risk factor *X*_*k*_, requiring at least five genome wide significant genetic instruments, assigning *a*_*k*_ *= 0* for factors with a lower number of instruments or non-significant causal effects (Bonferroni-corrected P < 0.05). MVMR advances this by regressing the exposure effect size matrix *B*_*e*_ *∈R*^*S*×*K*^ on the outcome effect sizes *β*_*o*_, setting *a*_*k*_ to the resultant causal effect estimates. Additionally, we contrast DMR’s performance with its linear counterpart, DMR-LIN, adding linear disease risk prediction function *f*. As performance benchmarks, we also included supervised models trained directly on individual-level follow-up labels. We primarily compared DMR against a longitudinal reference model (LRM) using the same risk prediction function as DMR. Additionally, we conducted comparisons with ElasticNet, RandomForest, and XGBoost models. Hyperparameters for all models with access to individual-level follow-up labels were optimized using an inner 5-fold cross-validation procedure (**Supplementary Information**). In simulation scenarios, these models aimed to minimize the mean squared error, while in the type 2 diabetes study, the objective was to minimize the binary cross-entropy loss.

### Simulations

#### Dataset generation

In our simulation study, we used 26 blood traits from 309,865 unrelated Europeans in the UK Biobank as potential risk factors. We crafted scenarios where a subset of these traits exerted a causal influence on the health outcome, with individual risk contributions following a J-shaped function—initially remaining negligible until surpassing a certain threshold, beyond which they increased linearly. The health outcome was generated as the sum of a linear combination of these non-linearly transformed causal exposures, a horizontal pleiotropy effect, and Gaussian noise. The horizontal pleiotropy effect was simulated as a direct genetic contribution from a subset of the variants associated with the blood traits. We systematically varied key parameters, such as the number of causal risk factors, and the proportions of the outcome variance explained by the risk factors and the horizontal pleiotropy effect, respectively. For each simulation parameter configuration, we considered 10 repeat experiments, utilizing distinct random seeds. Detailed descriptions of our simulation approach can be found in the **Supplementary Information**.

#### Evaluation framework

Adhering to a two-sample framework, the data was divided evenly into exposure and outcome cohorts, with this consistent split maintained throughout all simulated scenarios. Through the instrument selection procedure, 2904 independent genetic instruments were identified in the exposure cohort. Using the outcome cohort, the effects of these genetic instruments on the outcome were estimated—this step substitutes the real data analysis process of obtaining instrument effects on the outcome from external GWAS results. We trained DMR using 80% of the exposure cohort and evaluated the risk prediction accuracy on the remaining 20%. Prediction accuracy was assessed using Spearman correlation between the predicted and simulated risk values. Within this evaluation framework we compare DMR against DMR-LIN, UVMR and MVMR. We also compared DMR with models trained on individual-level follow-up data, including the LRM, ElasticNet, RandomForest, and XGBoost models (**Supplementary Figure A2**). For these models, we considered inner 5-fold cross-validation for hyperparameter selection (grid of explored values in **Supplementary Information**). Standard errors for all metrics were calculated from the results of 10 repeat experiments. To ensure the calibration of the evaluation procedure, the Spearman correlation of all MR-based models was verified to be compatible with zero in simulations without causal links (**Supplementary Figure A1**).

### Diabetes risk predictions

#### Cohort definition

We utilized DMR to predict 5-year T2D risk from 37 established risk factors [23]. For the outcome genetic effects and standard errors, we considered the external T2D GWAS summary statistics from [22], which excluded the UKBB cohort. For the exposure cohort, we considered 218,665 unrelated Europeans from UKBB who did not have diabetes at the time of assessment. After matching variants across the two datasets and excluding palindromic variants, the instrument selection procedure described above identified 6,077 independent genetic instruments associated with at least one of the 37 traits. More info on the longitudinal cohort definition can be found in **Supplementary Information**.

#### Evaluation framework

We compared DMR with MR-based models (DMR-LIN, UVMR, and MVMR) and models with direct access to individual-level follow-up data (LRM, ElasticNet, RandomForest, and XGBoost; **Supplementary Information**). Additionally, we included a polygenic risk score (PRS) predictor, computed externally by [24] and made available in UKBB through field 26285, for comparison. To assess T2D risk prediction accuracy, we employed the Area Under the Receiver Operating Characteristic Curve (AUC), using actual 5-year T2D risk as labels (derived from fields 41280 and 41270 using ICD10 code E11). To ensure robust significance testing and estimate standard errors, we conducted 50 repeat experiments with random 80%/20% splits for training and testing. Standard errors for all metrics were computed across these experiments, along with t-tests to assess performance improvements.

#### Interpretation of the learned T2D biomarker

To assess the risk predictor learned by DMR in the type 2 diabetes experiments, we employed analysis. Firstly, univariate associations were quantified using Spearman correlation between the risk predictor and each input exposure in out-of-sample individuals (**Supplementary Figure A4**). Secondly, to evaluate the model’s ability to capture nonlinear relationships among the selected risk factors, we visualized the learned contributions (normalized between 0 and 1) against observed values of these factors (**Figure 4c**). We observed that all reported values are highly consistent across all 50 repeat experiments.

### Imaging biomarkers of dementia

#### Cohort definition

We employed two-sample MR methods to estimate the 5-year AD risk based on brain T1 MRI features. For the exposure cohort, we selected 31,552 unrelated Europeans from UKBB with T1 brain imaging data. Out of the 153 brain volume features that are available in UKBB, we included 70 brain T1 MRI traits with at least 5 genetic instruments *P* < 5 ⋅ *10*^*−8*^ as risk factors in all MR models, for which we identified a total of 385 genetic instruments. More info on the exposure cohort can be found in **Supplementary Information**. For outcome genetic effects, we utilized external GWAS summary stats for AD in unrelated Europeans from [48].

#### Evaluation

We compared DMR with DMR-LIN, UVMR, and MVMR, and assessed AD risk prediction accuracy by AUC using actual 5-year AD risk as labels (derived from field 131036). Across 31,552 unrelated Europeans from UKBB with T1 brain imaging data, only 19 developed AD within 5 years. All models were trained on 80% of the healthy exposure cohort, while the remaining 20% along with 19 individuals with reported AD were used as a test set. To robustly test for significance and estimate standard errors, we conducted 50 repeat experiments, each employing different random 80%/20% splits.

#### Interpretation of the learned AD biomarker

For the Interpretation of the learned risk predictor we conducted a voxel-based association analysis using T1-weighted MRI scans that were registered to the MNI152 template [49–52]. Each voxel’s intensity was regressed against the out-of-sample individual’s risk predictor scores, adjusting for sex, age, UKB array type and the top 20 genetic principal components. This linear association test yielded p-values for contribution of the risk predictor to each voxel, which were transformed into a heatmap overlay on the MNI152 template using signed p-values *(− log*_*10*_ *(P)* ⋅*sign(β))*. Blue regions on the heat map indicate areas of volume decrease associated with increased risk predictor values, highlighting potential areas that correlate with higher AD risk (**Figure 4b**). Furthermore, we quantified the Spearman correlation between the risk predictor and each input imaging trait within the held-out validation set (**Figure 4c; Supplementary Figure A5**). Overall, both analyses displayed remarkable consistency across all experimental repeats, underscoring the reliability of our findings.

#### Use of Artificial Intelligence

In the preparation of this manuscript, we utilized the large language model GPT-4 (https://chat.openai.com/) for editing assistance, including language polishing and clarification of text. While this tool assisted in refining the manuscript’s language it was not used to generate contributions to the original research, data analysis, or interpretation of results. All final content decisions and responsibilities rest with the authors.

## Data access

No primary data were generated for this study. The UK Biobank dataset is available at https://biobank.ndph.ox.ac.uk/ukb/ after a registration and approval process. The type 2 diabetes GWAS summary statistics is available at http://www.type2diabetesgenetics.org/. The Alzheimer’s disease GWAS summary statistics is available at https://ctg.cncr.nl/software/summary_statistics.

## Declarations

### Competing interests

The authors declare that they have no competing interests.

## Acknowledgements

We would like to thank Julien Gagneur for feedback on the manuscript and providing resources. This research has been conducted using the UK Biobank Resource (Application Number 87065). FPC and DS were funded by the Free State of Bavaria’s Hightech Agenda through the Institute of AI for Health (AIH). LS acknowledges the support of the Friedrich-Alexander-Universität Erlangen-Nürnberg under the joint research school Munich School for Data Science (MUDS).

## Authors’ contributions

FPC conceived the study and supervised the work. DS, LG, FPC implemented the methods. DS, LG, LS, FPC analyzed the data. DS, LG, LS, FPC interpreted the results. DS, LG, LS, FPC wrote the paper.

## Notes

### Competing Interest Statement

The authors have declared no competing interest.

