## Supplementary_Information for "Genetics-driven Risk Predictions with Differentiable Mendelian Randomization"

### Supplementary Materials for "Genetics-driven Risk Predictions with Differentiable Mendelian Randomization"

**A2 Supplementary Figures** **9**

#### A1 Supplementary Information

##### A1.1 Related methods

**Risk predictions using longitudinal data.** Risk prediction from electronic health records traditionally employs longitudinal data [15, 33]. Predictive models used for this task range from simple methods combining questionnaire responses [25, 31] to ML supervised models integrating multiple data types [4]. To model event timing and censored data, survival prediction methods like Cox regression become essential [9, 16, 35]. In contrast to approaches using longitudinal data, DMR uses genetic effects on health outcomes as the supervisory signal to train risk predictors.

**MR methods.** The traditional MR methodology has seen numerous extensions tailored for varied settings. These encompass methods designed to address biases from weak instruments [41, 38, 40, 8, 39] and horizontal pleiotropy [7, 6, 21, 30, 42, 13, 37, 30, 41, 38, 27], as well as nonlinear models [11, 36, 34]. Among all these methods, DMR is most related to multivariable MR [32, 10], which performs concurrent analyses of multiple exposures under linear assumptions, and nonlinear MR [36, 11], which enables the investigation of how a single exposure's causal effect varies for different values of the exposure [36, 11]. Importantly, DMR extends these methods, enabling for the first time the learning of multivariable nonlinear functions for enhanced risk predictions.

**Machine Learning in Instrument Variable Analysis.** In the broader class of instrument variable analysis methods, the integration of machine learning (ML) techniques has recently begun to gain traction. For example, DeepIV leverages deep learning to predict treatment from high-dimensional instruments or intricate instrument structures [20, 3, 18]. In the realm of MR, Malina et al [26] combined MR with in silico mutagenesis to examine causal relationships between genomic marks and outcomes. Distinctly from these approaches, DMR leverages ML to learn nonlinear functions of multiple risk factors as aggregate exposures within the MR framework.

**Polygenic Risk Scores.** Historically, predicting disease risk using genetic data has been realized through polygenic risk scores (PRSs), which estimate individuals' genetic risk aggregating effects from associated genetic variants. This is commonly achieved through a GWAS, followed by procedures accounting for linkage disequilibrium to select independent loci [17, 29]. Notably, PRS can offer substantial potential for risk assessment and personalized healthcare [24, 5, 2], especially when integrated with risk trait data [24, 1, 22, 19, 23]. Unlike PRSs, which predict disease risk directly from genetic variants, DMR uses genetic data to learn risk predictors that are functions of multiple risk factors.

#### A1.2 Detailed Description of the DMR Model

For an exposure cohort consisting of $N$ individuals, $K$ risk factors, $C$ covariates, and $S$ independent genetic instruments associated with at least one of the $K$ risk factors, we define the following matrices and vectors:

- $\boldsymbol{X}$ represents the $N \times K$ matrix of risk factors;
- $\boldsymbol{F}$ represents the $N \times C$ matrix of covariates;
- $\boldsymbol{G}$ denotes the $N \times S$ matrix of genetic instruments;
- $\boldsymbol{\beta}_o$ is the $S \times 1$ vector of genetic effects on the disease outcome, obtained from external GWAS summary statistics;
- $\boldsymbol{s}_o$ is the $S \times 1$ vector of standard errors associated with $\boldsymbol{\beta}_o$, also obtained from external GWAS summary statistics.

Within the DMR framework, a differentiable function $f_{\boldsymbol{\phi}}$, parameterized by $\boldsymbol{\phi}$, transforms the input risk factors $\boldsymbol{X}$ into a single $N \times 1$ aggregate risk factor vector $\boldsymbol{e}(\boldsymbol{\phi}) = f_{\boldsymbol{\phi}}(\boldsymbol{X})$.* The DMR model assumes the aggregate risk factor $\boldsymbol{e}(\boldsymbol{\phi})$ as the exposure in an IVW regression model, formulated as:

$$\boldsymbol{\beta}_o \sim \mathcal{N}\left(\boldsymbol{\beta}_e(\boldsymbol{\phi})\alpha, \sigma^2 \operatorname{diag}(\boldsymbol{s}_o^2)\right). \tag{A.1}$$

*This operation is applied row-wise to each individual in $\boldsymbol{X}$.

Here, $\boldsymbol{\beta}_e(\boldsymbol{\phi})$ represents the marginal regression weights for each genetic variant in $\boldsymbol{G}$ against $\boldsymbol{e}(\boldsymbol{\phi})$, adjusting for covariates $\boldsymbol{F}$. For a single $N \times 1$ genetic instrument $\boldsymbol{g}$, the marginal effect can be computed through the following linear operator on $\boldsymbol{e}(\boldsymbol{\phi})$:

$$\underbrace{\mathcal{P}_0\left([\boldsymbol{g},\boldsymbol{F}]^T[\boldsymbol{g},\boldsymbol{F}]\right)^{-1}[\boldsymbol{g},\boldsymbol{F}]^T}_{\mathcal{M}(\boldsymbol{g},\boldsymbol{F})}\boldsymbol{e}(\boldsymbol{\phi}), \tag{A.2}$$

where $\mathcal{P}_0$ is a projection extracting the first element of the $(1+K)\times 1$ vector, which corresponds to the estimate of the effect size of $\boldsymbol{g}$. Thus, across all instruments, we have:

$$\boldsymbol{\beta}_e(\boldsymbol{\phi}) = h(\boldsymbol{e}(\boldsymbol{\phi}),\boldsymbol{G},\boldsymbol{F}) = \begin{bmatrix}\mathcal{M}(\boldsymbol{G}_{:,1},\boldsymbol{F})\boldsymbol{e}(\boldsymbol{\phi})\\ \cdots \\ \mathcal{M}(\boldsymbol{G}_{:,S},\boldsymbol{F})\boldsymbol{e}(\boldsymbol{\phi})\end{bmatrix}, \tag{A.3}$$

where $\boldsymbol{G}_{:,s}$ denotes the $s$-th column of $\boldsymbol{G}$, corresponding to the genotype vector of instrument $s$. For the functional form of $f_{\boldsymbol{\phi}}$, we adopt:

$$f_{\boldsymbol{\phi}}(\boldsymbol{x}) = \sum_{k=1}^{K} a_k \times \mathrm{ELU}(b_k x_k + c_k), \tag{A.4}$$

where the parameters $\boldsymbol{\phi} = \{a_1,\ldots,a_K,b_1,\ldots,b_K,c_1,\ldots,c_K\}$, and $\mathrm{ELU}(\cdot)$ is the Exponential Linear Unit activation function.

#### A1.3 Reduction to Multivariable MR for linear function

When adopting a linear function, DMR reduces to Multivariable MR. Indeed, when $e(\boldsymbol{\phi}) = \boldsymbol{X}\boldsymbol{\phi}$, Eq (A.3) becomes

$$\boldsymbol{\beta}_e(\boldsymbol{\phi}) = \begin{bmatrix}\mathcal{M}(\boldsymbol{G}_{:,1},\boldsymbol{F})\boldsymbol{X}\boldsymbol{\phi}\\ \cdots \\ \mathcal{M}(\boldsymbol{G}_{:,S},\boldsymbol{F})\boldsymbol{X}\boldsymbol{\phi}\end{bmatrix} = \underbrace{\begin{bmatrix}\mathcal{M}(\boldsymbol{G}_{:,1},\boldsymbol{F})\boldsymbol{X}\\ \cdots \\ \mathcal{M}(\boldsymbol{G}_{:,S},\boldsymbol{F})\boldsymbol{X}\end{bmatrix}}_{\boldsymbol{B}_e}\boldsymbol{\phi}, \tag{A.5}$$

where $\boldsymbol{B}_e$ are the $S \times K$ marginal effects of the $S$ genetic instruments on the $K$ traits. Replacing in Eq (A.8) we have

$$\boldsymbol{\beta}_o \sim \mathcal{N}\left(\boldsymbol{B}_e\underbrace{\boldsymbol{\phi}\alpha}_{\boldsymbol{\alpha}}, \sigma^2\mathrm{diag}(\boldsymbol{s}_o^2)\right), \tag{A.6}$$

where we redefined risk factor effects $\boldsymbol{\alpha}$. This is equivalent to the multivariable MR model introduced in [32, 10].

#### A1.4 Bayesian Extension

First, we note that replacing the parametric for of $f$ in Eq (A.4) in Eq (A.3), we have

$$\boldsymbol{\beta}_e(\boldsymbol{\phi}) = \underbrace{\begin{bmatrix}\mathcal{M}(\boldsymbol{G}_{:,1},\boldsymbol{F})\tilde{\boldsymbol{X}}(\boldsymbol{b},\boldsymbol{c})\\ \cdots \\ \mathcal{M}(\boldsymbol{G}_{:,S},\boldsymbol{F})\tilde{\boldsymbol{X}}(\boldsymbol{b},\boldsymbol{c})\end{bmatrix}}_{\boldsymbol{B}_e(\boldsymbol{b},\boldsymbol{c})}\boldsymbol{a}, \tag{A.7}$$

where $\boldsymbol{\phi} = \{\boldsymbol{a}, \boldsymbol{b}, \boldsymbol{c}\}$, and we introduced transformed risk factors $\tilde{\boldsymbol{X}}(\boldsymbol{b}, \boldsymbol{c})$ through the elu function, and the $S \times K$ marginal effects $\boldsymbol{B}_e(\boldsymbol{b}, \boldsymbol{c})$ of the $S$ genetic instruments on these $K$ transformed factors. Replacing this expression into the IVW model in Eq (A.8), we have:

$$\boldsymbol{\beta}_o \sim \mathcal{N}\left(\boldsymbol{B}_e(\boldsymbol{b}, \boldsymbol{c})\, \boldsymbol{a}, \sigma^2 \mathrm{diag}(\boldsymbol{s}_o^2)\right). \tag{A.8}$$

where we absorbed parameter $\alpha$ into $\boldsymbol{a}$. Next, we extend this model to a Bayesian setting to enhance robustness when the number of genetic instruments $S$ is limited. Specifically, we introduce priors over the model parameters $\boldsymbol{a}$, $\boldsymbol{b}$ and $\boldsymbol{c}$:

$$\boldsymbol{a} \sim \mathcal{N}(\boldsymbol{0}, \sigma_a^2 \boldsymbol{I}), \tag{A.9}$$
$$\boldsymbol{b} \sim \mathcal{N}(\boldsymbol{0}, \sigma_b^2 \boldsymbol{I}), \tag{A.10}$$
$$\boldsymbol{c} \sim \mathcal{N}(\boldsymbol{0}, \sigma_c^2 \boldsymbol{I}), \tag{A.11}$$

where $\sigma_a^2$, $\sigma_b^2$ and $\sigma_c^2$ are the variances of the priors. The log marginal likelihood of the model, integrating out the parameters, is given by:

$$\begin{aligned} p(\boldsymbol{\beta}_o|\sigma_a^2, \sigma_b^2, \sigma_c^2) &= \int \log \mathcal{N}\left(\boldsymbol{\beta}_o|\boldsymbol{B}_e(\boldsymbol{b}, \boldsymbol{c})\boldsymbol{a}, \sigma^2 \mathrm{diag}(\boldsymbol{s}_o^2)\right) \\ &\quad \times \mathcal{N}(\boldsymbol{a}|\boldsymbol{0}, \sigma_a^2\boldsymbol{I})\mathcal{N}(\boldsymbol{b}|\boldsymbol{0}, \sigma_b^2\boldsymbol{I})\mathcal{N}(\boldsymbol{c}|\boldsymbol{0}, \sigma_c^2\boldsymbol{I})\, d\boldsymbol{a}\, d\boldsymbol{b}\, d\boldsymbol{c} \\ &= \int \underbrace{\log \mathcal{N}\left(\boldsymbol{\beta}_o|\boldsymbol{0}, \sigma_a^2 \boldsymbol{B}_e(\boldsymbol{b}, \boldsymbol{c})\boldsymbol{B}_e(\boldsymbol{b}, \boldsymbol{c})^T + \sigma^2 \mathrm{diag}(\boldsymbol{s}_o^2)\right)}_{\log p(\boldsymbol{\beta}_o|\boldsymbol{b}, \boldsymbol{c}, \sigma_a^2)} \\ &\quad \times \mathcal{N}(\boldsymbol{b}|\boldsymbol{0}, \sigma_b^2\boldsymbol{I})\mathcal{N}(\boldsymbol{c}|\boldsymbol{0}, \sigma_c^2\boldsymbol{I})\, d\boldsymbol{b}\, d\boldsymbol{c}, \end{aligned} \tag{A.12}$$

where in the last passage we analytically integrated over $\boldsymbol{a}$. As the remaining integral is intractable, we employ mean-field variational inference to derive an Evidence Lower Bound (ELBO):

$$\begin{aligned} \mathrm{ELBO}(\sigma_a^2, \sigma_b^2, \sigma_c^2, \boldsymbol{\lambda}) &= \mathbb{E}_{q_{\boldsymbol{\lambda}}(\boldsymbol{b},\boldsymbol{c})}[\log p(\boldsymbol{\beta}_o|\boldsymbol{b}, \boldsymbol{c}, \sigma_a^2)] \\ &\quad - \mathrm{KL}[q_{\boldsymbol{\lambda}}(\boldsymbol{b}, \boldsymbol{c})||\mathcal{N}(\boldsymbol{b}|\boldsymbol{0}, \sigma_b^2\boldsymbol{I})\mathcal{N}(\boldsymbol{c}|\boldsymbol{0}, \sigma_c^2\boldsymbol{I})], \end{aligned} \tag{A.13}$$

where $q_{\boldsymbol{\lambda}}(\boldsymbol{b}, \boldsymbol{c})$ is the family of fully factorized Gaussian distributions on $\boldsymbol{b}$ and $\boldsymbol{c}$ parametrized by variational parameters $\boldsymbol{\lambda} = \{\boldsymbol{b}_m, \boldsymbol{b}_s, \boldsymbol{c}_m, \boldsymbol{c}_s\}$, and KL denotes the Kullback-Leibler divergence. The ELBO is maximized with respect to $\sigma_a^2$ and $\boldsymbol{\lambda}$ using gradient ascent, employing the reparametrization trick for gradients through the expectation term. Prior parameters $\sigma_b^2$ are $\sigma_c^2$ are set to 0.8 and 3 respectively, based on our prior assumptions on the type of nonlinearities we will see in the data.

#### A1.5 Standard ML model baselines

In this section, we elaborate on the training procedure and implementation details of the supervised machine learning models that have access to individual-level follow-up labels: ElasticNet (EN), RandomForest (RF), and XGBoost (XGB).

**Training Procedure.** Hyperparameter optimization was conducted for all models using a grid search approach. For each combination of parameters within the grid, an inner 5-fold cross-validation was performed. The performance of each hyperparameter configuration was assessed based on the average metric across all folds. After identifying the optimal hyperparameters, the models were refitted on the entire training dataset and subsequently evaluated on the same held-out data as used for the DMR model. In the simulations, the mean squared error served as the performance metric for hyperparameter tuning, while in the type 2 diabetes (T2D) experiments, binary cross-entropy was used for the RandomForest and XGBoost models.

**Implementation Details.** The ElasticNet and RandomForest models were implemented using the regressor and classifier classes, respectively, from the `scikit-learn` [28] Python package for simulations and T2D experiments. For XGBoost, we employed the respective regressor and classifier implementations from the `XGBoost` Python package [12]. The following hyperparameter grids were utilized for optimization across both simulations and T2D experiments:

**ElasticNet.** The hyperparameter grid for ElasticNet was as follows:

- `l1_ratio`: [0.1, 0.5, 0.7, 0.9, 0.95, 0.99, 1.0]
- `eps`: 5.0e-4
- `n_alphas`: 500
- `max_iter`: 5000

**RandomForest.** The hyperparameter grid for RandomForest included:

- `n_estimators`: [100, 200, 300]
- `max_depth`: [10, 20, 30, None]
- `min_samples_split`: [2, 5, 10]
- `min_samples_leaf`: [1, 2, 4]

**XGBoost.** The hyperparameter grid for XGBoost was composed of:

- `n_estimators`: [100, 200, 300]
- `max_depth`: [3, 9, 0]
- `learning_rate`: [0.01, 0.05, 0.1]
- `subsample`: [0.6, 0.8, 1.0]
- `colsample_bytree`: [0.6, 0.8, 1.0]

#### A1.6 Simulations design

For $N$ individuals, $K$ exposures, and $S$ independent genetic instruments, let $\boldsymbol{E}$ denote the $N \times K$ exposure matrix, and $\boldsymbol{G}$ denote the $N \times S$ genetic instruments matrix. The outcome is simulated as the sum of three contributions:

$$\boldsymbol{y} = \underbrace{\boldsymbol{e}_{\text{sim}}}_{\text{simulated risk}} + \underbrace{\boldsymbol{h}_{\text{pleio}}}_{\text{horizontal pleiotropy}} + \underbrace{\boldsymbol{\epsilon}}_{\text{Gaussian noise}} \tag{A.14}$$

Given the number of causal exposures ($K_c$), the variance explained by simulated risk ($v_{\text{caus}}$), and the variance explained by horizontal pleiotropy ($v_{\text{pleio}}$), each term was simulated as described in the following:

1. **Simulated risk** ($\boldsymbol{e}_{\text{sim}}$)
   - Select $K_c$ causal exposures with indices $\{i_1, \dots, i_{K_c}\}$
   - Simulate risk

$$\boldsymbol{e}_{\text{sim}} = \frac{1}{K_c} * \sum_{k}^{K_c} \text{std}(\text{ELU}(\boldsymbol{E}_{:,i_k} - q(\boldsymbol{E}_{:,i_k}, 0.75))) \tag{A.15}$$

   where ELU denotes the Exponential Linear Unit function and std is the standardization function across $N$ individuals. This procedure ensures that each causal exposure explains the same variance
   - Rescale such that $\boldsymbol{e}_{\text{sim}}$ has variance $v_{\text{caus}}$

$$\boldsymbol{e}_{\text{sim}} \leftarrow \sqrt{v_{\text{caus}}}\, \boldsymbol{e}_{\text{sim}} \tag{A.16}$$

2. **Horizontal Pleiotropy** ($\boldsymbol{h}_{\text{pleio}}$)
   - Select 10% causal SNPs and standardize variants, $\boldsymbol{G}_c = \text{std}(\boldsymbol{G}[:, \{s_1, \dots, s_H\}])$
   - Simulate genetic effects and standardize

$$\boldsymbol{h}_{\text{pleio}} = \text{std}(\boldsymbol{G}_c \boldsymbol{b}), \quad \text{with } \boldsymbol{b} \overset{\text{iid}}{\sim} \text{Unif}(\{-1, 1\}) \tag{A.17}$$

   - Rescale such that $\boldsymbol{h}_{\text{pleio}}$ has variance $v_{\text{pleio}}$

$$\boldsymbol{h}_{\text{pleio}} \leftarrow \sqrt{v_{\text{pleio}}}\, \boldsymbol{h}_{\text{pleio}} \tag{A.18}$$

3. **Gaussian noise** ($\boldsymbol{\epsilon}$)

$$\boldsymbol{\epsilon} = \sqrt{1 - v_{\text{caus}} - v_{\text{pleio}}} \cdot \text{std}(\boldsymbol{\eta}), \quad \text{with } \boldsymbol{\eta} \overset{\text{iid}}{\sim} \mathcal{N}(0, 1) \tag{A.19}$$

#### A1.7 Definition of the cohort for T2D onset prediction

**Cohort definition.** To test the performance of DMR for the prediction of T2D development, we retrospectively created a cohort from UKBB (see Figure 4a). We considered unrelated Europeans from UKBB who did not have diabetes at the time of assessment. This was ensured by filtering the population based on the following criteria:

- Do not have reports of any kind of diabetes before the baseline or within the first six months after the assessment. The latter guarantees that the diagnosis was not given directly after, based on the data collected at the baseline. The comparison is done using the following fields: 130706, 130708, 130710, 130712, 130714, 130716, 130718, and Instance 0.0 of field 53.
- Do not have a critical level of the main biomarker for T2D, HbA1c (HbA1c $<$ 48 mmol/mol) at the baseline (Instance 0.0, field 30750).
- Medication-based exclusion of patients from the initial cohort that reported taking one of the following drugs at the time of baseline assessment: 'metformin', 'glibenclamide', 'glimepiride', 'repaglinide', 'nateglinide', 'troglitazone', 'pioglitazone', 'rosiglitazone', 'rosiglitazone 1mg/metformin 500mg tablet', 'acarbose', 'gliclazide', 'glipizide product', 'tolbutamide', 'insulin product'. The data was taken from field 20003.

As we were aiming to predict the T2D development in 5 years after assessment, some additional censoring was also implemented. Patients who have died or did not have Health and Social Care Information Centre (HES) or General Practitioner (GP) records during the 5 years of follow-up would not have received the right T2D diagnosis if they had the disease. Hence, they were removed from the cohort.
After these exclusions, the size of the cohort was 218,665 people. According to the ICD10 medical records in fields 41270 and 41280, 1,490 patients in the cohort developed type 2 diabetes (code E11) within the next 5 years and were further considered as "cases". The rest of the filtered cohort was considered to be controls.

**Risk factor selection.** The prediction of T2D is based on the following 37 predictive factors:

- 26 blood biomarkers (4 were excluded because of high percentage of missing data):
  - Albumin (field 30600)
  - Alkaline Phosphatase (field 30610)
  - Alanine Aminotransferase (field 30620)
  - Apolipoprotein A (field 30630)
  - Apolipoprotein B (field 30640)
  - Aspartate Aminotransferase (field 30650)
  - Urea (field 30670)
  - Calcium (field 30680)
  - Cholesterol (field 30690)
  - Creatinine (field 30700)
  - C-Reactive Protein (field 30710)
  - Cystatin C (field 30720)
  - Gamma-Glutamyltransferase (field 30730)
  - Glucose (field 30740)
  - Glycated Hemoglobin (field 30750)
  - HDL Cholesterol (field 30760)
  - IGF-1 (field 30770)
  - LDL Direct (field 30780)
  - Phosphate (field 30810)
  - SHBG (field 30830)
  - Total Bilirubin (field 30840)
  - Testosterone (field 30850)
  - Total Protein (field 30860)
  - Triglycerides (field 30870)
  - Urate (field 30880)
  - Vitamin D (field 30890)
- Diastolic blood pressure, automated reading (field 4079)

- Systolic blood pressure, automated reading (field 4080)
- Pulse rate, automated reading (field 102)
- Waist-Hips circumference ratio, calculated from Waist circumference (field 48) and Hip circumference (field 49) is
- BMI, Body mass index (field 21001)
- 6 blood-count features (easiest to measure):
  - Hemoglobin concentration (field 30020)
  - Mean corpuscular volume (field 30040)
  - Mean platelet (thrombocyte) volume (field 30100)
  - Red blood cell (erythrocyte) count (field 30010)
  - White blood cell (leukocyte) count (field 30000)
  - Platelet count (field 30080)

Missing values within the data were imputed using the mean of the respective feature.

#### A1.8 Definition of the cohort for Alzheimer's Dementia prediction

To test the performance of DMR for the prediction of dementia, we created a cohort from UKBB with 31,552 unrelated individuals of European descent with available T1 MRI brain imaging data. As predictive features, we used brain T1 MRI volume measurements from categories 1101 and 1102 of UKBB. Out of these 153 brain volume features, we selected 70 traits, which had at least 5 genetic instruments ($P < 5 \cdot 10^{-8}$) identified by our IV selection procedure. Missing values within the data were imputed using the mean of the respective feature.

#### A2 Supplementary Figures

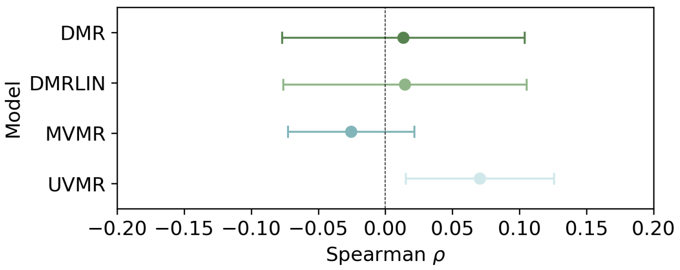

Figure A1: **Model calibration under simulations with no causal effects.** The figure presents Spearman correlation coefficients $\rho$ between simulated risk factors and predictions from DMR, DMR-LIN, UVMR, MVMR, and LRM on the held-out validation set, when no causal effects are simulated. Error bars denote standard errors across 10 repeat experiments. The dashed line at $\rho = 0$ indicates the expected random chance correlations in this simulated setting. These results confirm calibration of our simulation and evaluation procedure.

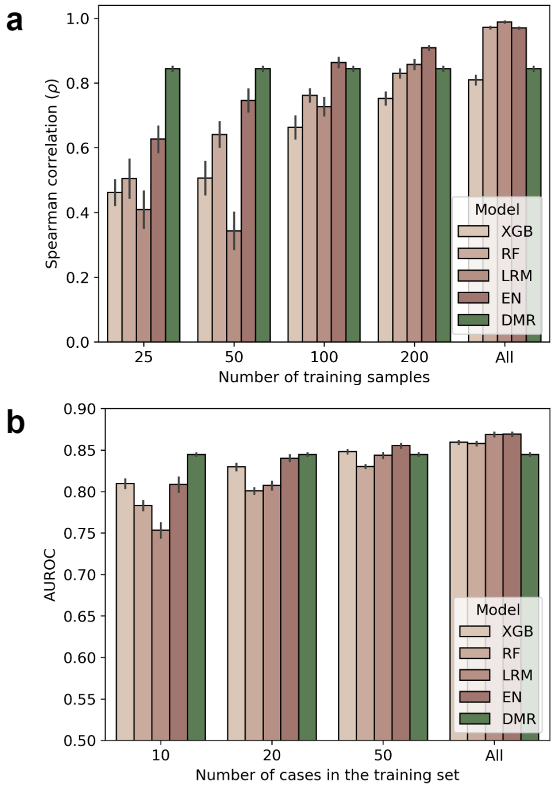

Figure A2: **Comparison with supervised ML models with access to individual-level follow-up labels.** Panel (**a**) showcases the Spearman correlation across different training set sizes in data simulations, comparing DMR to supervised ML models with access to individual-level follow-up labels. For this comparison, we considered ElasticNet (EN), RandomForest (RF), XGBoost (XGB), and small neural network adopting same risk prediction funciton of DMR (Longitudinal Reference Model; LRM). Panel (**b**) shows the area under the receiver operating characteristic curve (AUROC) for 5-year T2D risk predictions comparing DMR with the same set of models. Here the models were trained by restricting the number of cases in the training set, where controls are sampled such that the ration between cases and controls remains consistent for the different numbers shown in the plot.

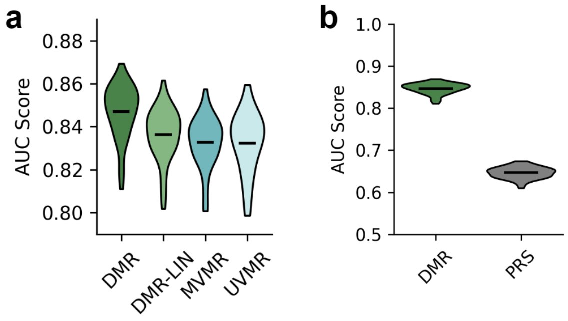

Figure A3: **Comparison of AUC Scores Across Risk Prediction Models.** Panel (**a**) displays the distribution of AUC scores for DMR, DMR-LIN, MVMR, and UVMR models. Panel (**b**) contrasts the distribution of the AUC score of DMR against a polygenic risk score (PRS) model.

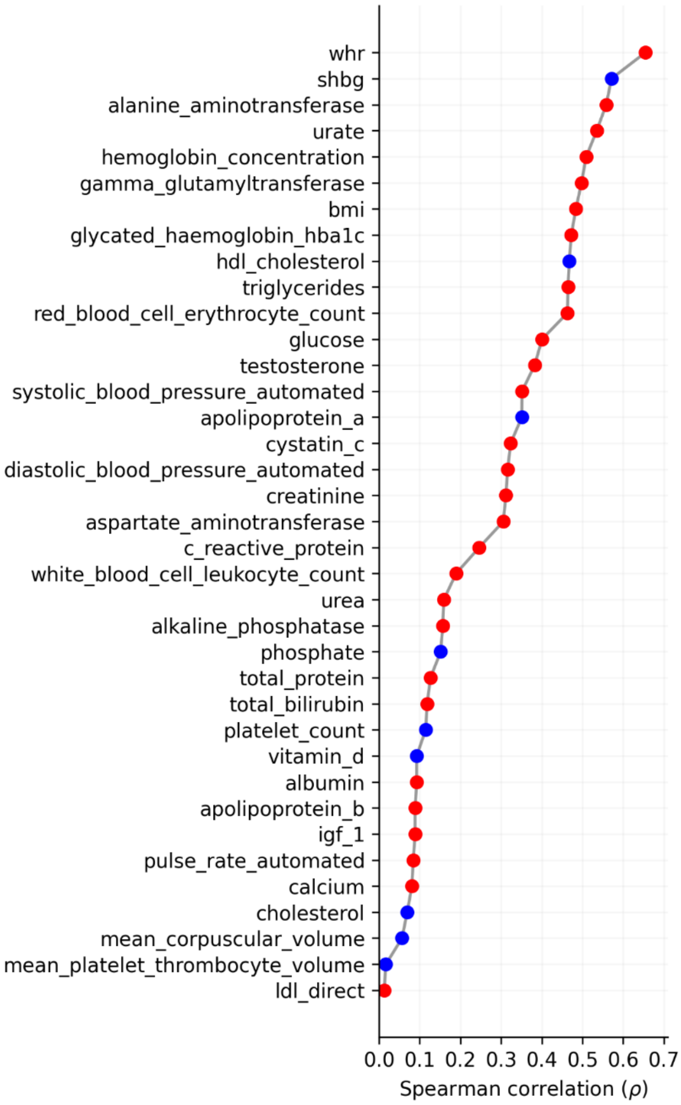

Figure A4: **Correlation of T2D risk predictor with clinical blood markers.** Shown are the Spearman correlation coefficients ($\rho$) between the learned risk predictor of DMR and the analyzed risk factors. Red dots represent markers with positive correlations, while blue dots indicate negative correlations.

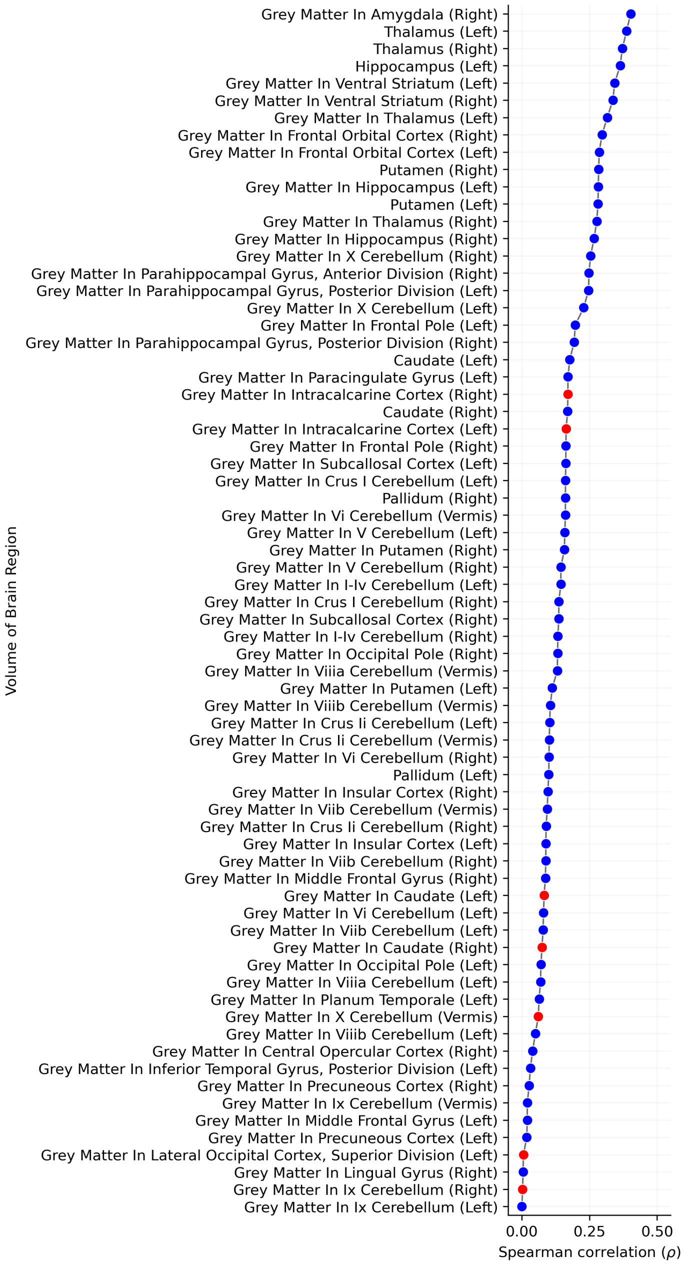

Figure A5: **Correlation of AD risk predictor with T1 imaging traits.** Shown are the Spearman correlation coefficients ($\rho$) between the DMR risk predictor for AD and the analyzed T1 MRI features. Red dots represent markers with positive correlations, while blue dots indicate negative correlations. The strongest negative correlations, indicative of gray matter reduction, are observed in regions known to be impacted by Alzheimer's pathology, such as the amygdala and hippocampus [14].

[2] Adebowale Adeyemo, Mary K. Balaconis, Deanna R. Darnes, Segun Fatumo, Palmira Granados Moreno, Chani J. Hodonsky, Michael Inouye, Masahiro Kanai, Kazuto Kato, Bartha M. Knoppers, Anna C. F. Lewis, Alicia R. Martin, Mark I. McCarthy, Michelle N. Meyer, Yukinori Okada, J. Brent Richards, Lucas Richter, Samuli Ripatti, Charles N. Rotimi, Saskia C. Sanderson, Amy C. Sturm, Ricardo A. Verdugo, Elisabeth Widen, Cristen J. Willer, Genevieve L. Wojcik, and Alicia Zhou. Responsible use of polygenic risk scores in the clinic: potential benefits, risks and gaps. Nature Medicine, 27(11):1876–1884, 2021.

[3] Andrew Bennett, Nathan Kallus, and Tobias Schnabel. Deep generalized method of moments for instrumental variable analysis. Advances in neural information processing systems, 32, 2019.

[19] Seok-Ju Hahn, Suhyeon Kim, Young Sik Choi, Junghye Lee, and Jihun Kang. Prediction of type 2 diabetes using genome-wide polygenic risk score and metabolic profiles: A machine learning analysis of population-based 10-year prospective cohort study. EBioMedicine, 86, 2022.

[20] Jason Hartford, Greg Lewis, Kevin Leyton-Brown, and Matt Taddy. Deep iv: A flexible approach for counterfactual prediction. In International Conference on Machine Learning, pages 1414–1423. PMLR, 2017.

[21] Fernando Pires Hartwig, George Davey Smith, and Jack Bowden. Robust inference in summary data mendelian randomization via the zero modal pleiotropy assumption. International journal of epidemiology, 46(6):1985–1998, 2017.

[22] Michael Inouye, Gad Abraham, Christopher P Nelson, Angela M Wood, Michael J Sweeting, Frank Dudbridge, Florence Y Lai, Stephen Kaptoge, Marta Brozynska, Tingting Wang, et al. Genomic risk prediction of coronary artery disease in 480,000 adults: implications for primary prevention. Journal of the American College of Cardiology, 72(16):1883–1893, 2018.

[23] Kristi Läll, Reedik Mägi, Andrew Morris, Andres Metspalu, and Krista Fischer. Personalized risk prediction for type 2 diabetes: the potential of genetic risk scores. Genetics in Medicine, 19(3):322–329, 2017.

[24] Samuel A Lambert, Gad Abraham, and Michael Inouye. Towards clinical utility of polygenic risk scores. Human molecular genetics, 28(R2):R133–R142, 2019.

[33] Maren E Shipe, Stephen A Deppen, Farhood Farjah, and Eric L Grogan. Developing prediction models for clinical use using logistic regression: an overview. Journal of thoracic disease, 11(Suppl 4):S574, 2019.

[41] Qingyuan Zhao, Jingshu Wang, Gibran Hemani, Jack Bowden, and Dylan S Small. Statistical inference in two-sample summary-data mendelian randomization using robust adjusted profile score. 2020.

[42] Zhihong Zhu, Zhili Zheng, Futao Zhang, Yang Wu, Maciej Trzaskowski, Robert Maier, Matthew R Robinson, John J McGrath, Peter M Visscher, Naomi R Wray, et al. Causal associations between risk factors and common diseases inferred from gwas summary data. Nature communications, 9(1):1–12, 2018.
